# pLAST - a tool for rapid comparison and classification of bacterial plasmid sequences

**DOI:** 10.1101/2025.11.27.689987

**Authors:** Kamil Krakowski, Małgorzata Orlowska, Kamil Kaminski, Dariusz Bartosik, Stanislaw Dunin-Horkawicz

## Abstract

**Motivation:** The increasing number of fully sequenced bacterial plasmids being annotated and cataloged has prompted the development of computational tools for comparing and classifying them. Existing approaches typically compare full-length DNA sequences (e.g., Mash, BLASTn) or translated open reading frames (ORFs) (e.g., BLASTp, DIAMOND), with plasmid-level scores obtained by aggregating ORF-to-ORF similarities; however, they are either restricted to closely related plasmids or become computationally demanding in large-scale analyses.

**Results:** We describe pLAST (plasmid Language Analysis and Search Tool), a plasmid-search tool built using word2vec representations capturing both ORF-to-ORF similarity and gene neighborhood conservation. Benchmarks indicate that pLAST outperforms both DNA- and ORF-based methods in identifying functionally similar plasmids and, compared to the widely used Mash, it achieves 37%, 30%, and 13% improvements in detecting shared mating-pair formation (MPF) system, relaxase, and *oriT* types, respectively. This performance scales to datasets comprising thousands of sequences, as exemplified by clustering analyses of ∼56,000 plasmids, which reveal expected functional groups. Beyond global similarity, pLAST also returns per-ORF plasmid-plasmid alignments, enabling detection of shared functional modules.

**Availability and implementation:** pLAST is freely accessible as a web server at https://plast.lbs.cent.uw.edu.pl/ and available as a Python module along with a precomputed database at https://github.com/labstructbioinf/pLAST for customized analysis.

## Introduction

Plasmids are extrachromosomal replicons that drive horizontal gene transfer, spreading antibiotic and metal resistance genes as well as virulence factors (Coluzzi and Rocha, 2025; Dewan and Uecker, 2023). Given their biological and evolutionary significance, dedicated plasmid databases are expanding rapidly. The PLSDB database has increased from ∼34k non-redundant entries in 2021 to ∼72k in the 2025 update (Molano *et al*., 2025). Non-dereplicated resources contain even more sequences, with IMG/PR and PlasmidScope databases hosting ∼700k and ∼852k plasmid sequences, respectively, including plasmids reconstructed from metagenomes and metatranscriptomes (Camargo *et al*., 2024; Li *et al*., 2025).

Searching these resources relies on tools that compare a query plasmid against database entries. Common options include DNA-based methods such as Mash (Ondov *et al*., 2016) and BLASTn (Altschul *et al*., 1990), as well as protein-based approaches leveraging ORF-to-ORF similarities returned by tools like BLASTp (Altschul *et al*., 1990) and DIAMOND (Buchfink *et al*., 2015). DNA methods typically compare whole plasmid nucleotide sequences to assess global similarity between plasmids (Camargo *et al*., 2024; Schmartz *et al*., 2021), whereas ORF-based frameworks aggregate ORF-to-ORF hits for gene-content comparison and module detection (Suzuki *et al*., 2020; Teixeira *et al*., 2023).

The application of Mash or BLASTn to plasmid comparison has key limitations: first, they operate on DNA sequences, which diverge faster than protein sequences, so for distant relationships, the homology signal often falls below detection; second, plasmids are modular and often mosaic, so shared sequence is frequently discontinuous, and accessory regions and mobile elements can distort global similarity measures (Redondo-Salvo *et al*., 2020); third, methods relying on sequence alignment (BLASTn) do not inherently account for sequence circularity, which can inflate estimated distances between otherwise highly similar plasmids, a problem explicitly targeted by some methods (Ayad and Pissis, 2017; Fernandes *et al*., 2009; Grossi *et al*., 2016).

These problems, highlighted in recent discussions on plasmid similarity metrics (Matlock *et al*., 2024), are partly mitigated by ORF-based methods, which are generally more robust for detecting distant homology. They do not explicitly model circularity or modular architectures, but comparing ORFs independently makes results largely insensitive to sequence rotation and gene order. However, the downside is that ignoring gene order discards synteny, an evolutionarily informative signal; moreover, protein-level matches that do not consider genomic context can be confounded by paralogs, especially those originating from recent gene duplications (Maida *et al*., 2014).

Here, we describe pLAST, a method that encodes ORF-to-ORF similarities into synteny-informed numerical representations, allowing functionally meaningful searches of plasmid databases and scalable clustering of large plasmid datasets.

## Methods and Implementation

### Dataset preparation

More than 56,000 plasmids were retrieved from the NCBI RefSeq nucleotide database (accessed April 2025) by querying it for plasmid records and restricting results to complete, circular sequences 5 kb to 10 Mb in length. The plasmids obtained were then annotated with MOB-suite (v3.1.9) (Robertson and Nash, 2018) and their protein sequences (>5.4 million in total) were clustered with MMseqs2 (*17*.*b804f*) (-s 7.5 --cluster-mode 1 --cov-mode 0 --min-seq-id 0.3 -c 0.8), yielding ∼386,000 unique clusters. In parallel, proteins were annotated using eggNOG-mapper (*2*.*1*.*12*; eggNOG DB *5*.*0*.*2*) (Huerta-Cepas *et al*., 2019; Cantalapiedra *et al*., 2021) and, where possible, assigned to the orthologous group (OG) with the lowest E-value, resulting in ∼64,000 unique OGs.

### Model training

Plasmids were encoded as sequences of ORFs, represented either by MMseqs2 cluster identifiers or by eggNOG orthologous group (OG) identifiers (yielding two pLAST variants). To avoid dependence on the arbitrary start coordinate in linearized RefSeq records, during dataset creation each ORF chain was randomly circularly permuted and, with 50% probability, reversed in order. Next, a grid of word2vec models was trained with Gensim (4.3.3) (Řehůřek and Sojka, 2010), covering two architectures (skip-gram and CBOW), vector dimensions of 64 or 128, five context window sizes (4, 8, 16, 32, 42), two frequency cutoffs (min-count = 1 or 2), and negative sampling (negative = 15). Each model was trained for 30 epochs, producing embeddings for every unique ORF class present after min-count filtering; the effective vocabulary sizes were 386,497 (MMseqs2, min-count = 1), 155,408 (MMseqs2, min-count = 2), 64,443 (eggNOG, min-count = 1), and 44,206 (eggNOG, min-count = 2).

For each model, a placeholder “blank-ORF” vector was computed by averaging all ORF embeddings. Subsequently, plasmid embeddings were computed as the weighted mean of their ORF embeddings, with ORF-specific weights given by corpus-wide inverse document frequency (rarer ORFs receive higher weight, common ORFs lower weight); ORFs missing a cluster assignment or whose assigned cluster identifier was missing in the given model received the placeholder embedding with a fixed weight of 0.01.

### Model evaluation

We evaluated how well plasmid embeddings from each model recover MOB-suite clustering at the primary and secondary levels, defined at Mash distance ≤ 0.06 and ≤ 0.025, respectively (Robertson *et al*., 2020). For each level, we constructed a balanced benchmark by randomly sampling 50 plasmids per MOB-suite cluster at that level (from clusters with ≥50 members). We then clustered the per-plasmid embeddings with K-means (k equal to the number of clusters at that level) and compared assignments to MOB-suite clusters using the Adjusted Rand Index (ARI). As a control, we also computed ARI for plasmid embeddings obtained by averaging ORF embeddings after randomly permuting the ORF-to-embedding mapping. For downstream analyses, we retained only models showing a positive ARI gain over their shuffled controls at both levels.

### Functional benchmark

In the benchmark, we focused on conjugative and mobilizable plasmids by selecting those with at least two of the following MOB-suite annotations: relaxase type, mating-pair formation (MPF) system type, or *oriT* type. Within each MOB-suite primary cluster of the resulting ∼23,000 plasmids, ∼10% were randomly sampled, ensuring at least one representative per MOB-suite secondary cluster. Primary clusters reduced to fewer than three plasmids after subsampling were excluded, yielding 1,772 plasmids from 159 primary clusters. One plasmid was then randomly selected per primary cluster to serve as a query (159 queries in total); the remaining plasmids formed the database (1,613 plasmids). The benchmark set is dominated by Enterobacteriaceae (84.1%), followed by Moraxellaceae (2.5%), Staphylococcaceae (1.7%), Bacillaceae (1.7%), and Enterococcaceae (1.6%). Together, these top five taxa comprise ∼91.6% of the sequences, reflecting the distribution of well-annotated RefSeq plasmids; the set remains highly non-redundant due to MOB-suite cluster-based subsampling.

The 159 queries were used to search the benchmark database with pLAST, where similarity was computed as the cosine between unit-normalized plasmid embeddings, and with four baselines: (i) BLASTp (E-value cut-off = 1e-3) and (ii) DIAMOND (E-value cut-off = 1e-3 and “--more-sensitive” option), both scored as the average of per-ORF best-hit bit scores over the query ORFs (before aggregation, greedy one-to-one matching of query and target ORFs within each plasmid pair and ordering hits by higher bit score and lower E-value was applied); (iii) Mash (k = 19, sketch size = 10^5), with similarity defined as 1 − distance; and (iv) BLASTn, with similarity defined as ANI×AF, where ANI is the average nucleotide identity over uniquely covered positions in the query, and AF (alignment fraction) is defined as *min*(*unique query coverage, unique subject coverage*) divided by the length of the shorter sequence, i.e. the fraction of the shorter sequence that is aligned in both sequences.

For each query, we computed the mean Jaccard index between its MOB-suite annotations and those of the top P% most similar database hits for that query (P samples the range 1 to 50%), then averaged these scores across queries. We performed this analysis separately for three annotation categories: relaxase type, mating-pair formation (MPF) system type, and origin of transfer (*oriT)* type.

Annotations marked as “unknown” in either the query or the hits were excluded. This yields performance curves and area-under-the-curve (AUC) scores (Figure 1), which we use to select the best-performing pLAST model and to compare it with BLASTp, DIAMOND, Mash, and BLASTn. For both the eggNOG- and MMseqs2-based variants, the top model uses CBOW with 64-dimensional vectors, a window size of 4, and min-count = 2.

**Figure 1.**
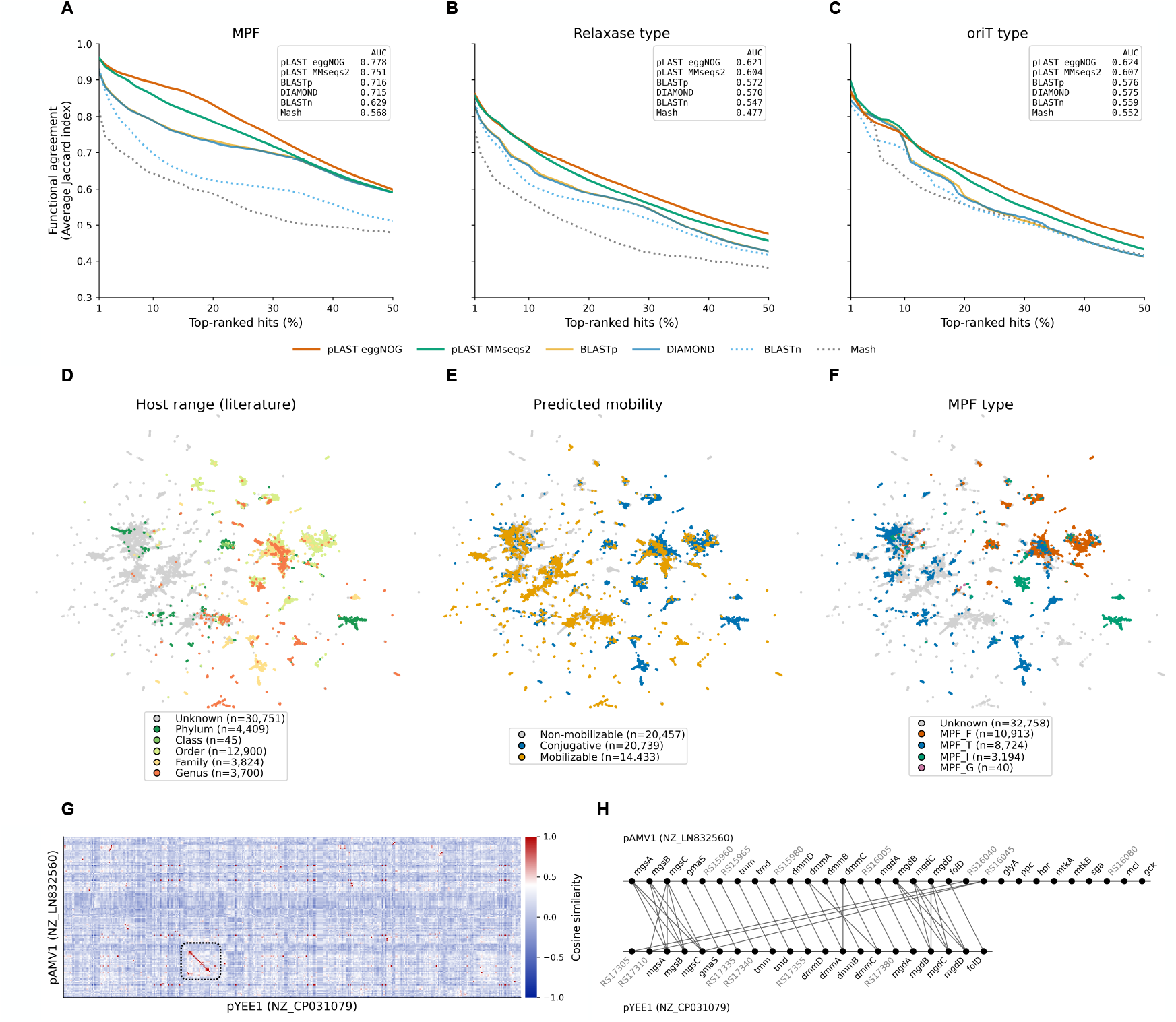
Performance of pLAST and other methods in detecting functionally similar plasmids. Three MOB-suite annotations are evaluated: (A) MPF (mating-pair formation) type, (B) relaxase type, and (C) *oriT* type. Solid lines denote protein-level methods (pLAST eggNOG, pLAST MMseqs2, BLASTp, DIAMOND); dotted lines denote nucleotide-level methods (BLASTn, Mash). The x-axis shows the percentage P of top-ranked database hits retained, and the y-axis shows the mean Jaccard index between each query’s annotations and those of its top P% hits, averaged across the 159 queries. AUC values are shown in each inset. Panels D-F display UMAP projections of pLAST MMseqs2 embeddings for ∼56,000 plasmids, colored by host range (D), predicted mobility (E), and MPF type (F) from MOB-suite. (G) Per-ORF cosine-similarity matrix between the plasmids pAMV1 (NZ_LN832560) and pYEE1 (NZ_CP031079). (H) Zoom-in on the dotted box in (G), highlighting a multi-gene syntenic region; dots are ORFs and connecting lines indicate ORF pairs with embedding cosine similarity ≥ 0.6.

## Results

pLAST was built by training word2vec models (originally developed for natural language processing) on plasmids encoded as sequences of ORFs, each represented by the identifier of the protein family it belongs to. These families were either annotated using the eggNOG database or derived by clustering ∼5.4 million translated ORFs from ∼56,000 plasmids with MMseqs2, yielding two model variants: pLAST eggNOG and pLAST MMseqs2 (see Methods for details). For each plasmid, with ORFs encoded as eggNOG or MMseqs2 families, the trained model computes a numerical representation (embedding) that captures patterns of ORF presence and short-range co-occurrence along the plasmid. These embeddings can then be quickly compared across plasmids to estimate their similarity.

To evaluate pLAST performance, we benchmarked it together with DNA sequence-based (BLASTn, Mash) and protein sequence-based (BLASTp, DIAMOND) methods for their ability to retrieve functionally similar plasmids. We constructed a representative set of ∼1,700 conjugative and mobilizable plasmids and assessed how well the similarity scores returned by each method agree with shared relaxase, *oriT*, and MPF types as defined by MOB-suite (Figure 1 A-C). Each of the three functional categories poses a distinct challenge: relaxase type is determined by a single locus, sometimes with a few accessory genes; MPF is a large, multi-gene module spanning many ORFs and often arranged in operons with mosaic variation; and *oriT* is a cis-acting DNA element that is not directly evaluable by protein-based methods.

Across all three functional categories, pLAST eggNOG and pLAST MMseqs2 achieved the best performance, followed by BLASTp and DIAMOND, then BLASTn, and Mash (Figure 1). In this benchmark, DNA-based methods underperform because conservation of protein-level functional modules is not well captured at the nucleotide level, neither by global k-mer sketch similarity (Mash) nor by local alignments (BLASTn). By contrast, protein-based methods perform substantially better, as they are more robust to sequence divergence and genomic rearrangements. The superior performance of pLAST comes from combining protein-level similarity (eggNOG or MMseqs2 families) with local co-occurrence context captured by the word2vec model. Notably, protein-based methods, including pLAST, also outperformed DNA-based methods in the *oriT* benchmark. Although *oriT* is a DNA element, these methods work well because *oriT* typically resides within conserved gene neighborhoods with characteristic protein composition. Moreover, relaxase type is often tightly coupled to its cognate *oriT*, so performance on the *oriT* class is partly reflected by performance on relaxase type.

In terms of runtime, searching the benchmark database with 159 queries took 15 minutes, 7 minutes, 3 minutes, and under one minute for BLASTp, BLASTn, DIAMOND, and Mash, respectively. By comparison, pLAST returns results within seconds because it operates on numerical representations that can be compared efficiently. Comparisons of these representations scale easily to much larger datasets, as exemplified by clustering ∼56,000 RefSeq plasmids, which recovers functionally coherent groups whose mobility gene repertoires are consistent with the reported host range (Figure 1 D-F). In practice, computing a numerical representation for a plasmid takes a few seconds with MMseqs2-GPU and about 1-2 minutes with eggNOG-mapper, depending on the number of ORFs.

The benchmark (Figure 1 A-C) and the clustering (D-F) described above rely on global plasmid-plasmid similarity scores; however, pLAST also returns per-ORF alignments, enabling the characterization of multi-gene regions shared between plasmids. For example, the methylotrophy islands (MEIs) are modular cassettes enabling the use of single-carbon (C1) compounds as a carbon and energy source. In the pAMV1 plasmid of *Paracoccus aminovorans* JCM 7685 (NZ_LN832560), the MEI comprises modules for substrate uptake and oxidation, THF-linked C1 processing, and assimilation via the serine cycle (Czarnecki *et al*., 2017). Its truncated variant, lacking the serine-cycle module (and thus C1 assimilation), has been reported in the *Paracoccus yeei* CCUG 32053 pYEE1 plasmid (NZ_CP031079; (Czarnecki and Bartosik, 2019)). The pLAST alignment of pAMV1 and pYEE1 (Figure 1 G-H) shows that, despite low overall similarity, they share the MEI region and reveals differences, including the loss of the serine-cycle module and a circular rearrangement that relocates two ORFs to the beginning of the cassette.

pLAST is available as a web server at https://plast.lbs.cent.uw.edu.pl/. The server enables searches against a precomputed RefSeq plasmid database using either eggNOG- or MMseqs2-based models. The resulting matches can be analysed globally via UMAP projections, such as those shown in Figure 1D–F, or individually by inspecting pairwise alignments. Together, these features enable rapid and comprehensive characterization of plasmids of interest in the context of the entire bacterial plasmidome, as well as assessment of the genetic diversity of these replicons and the rearrangements that give rise to their variants.

## Acknowledgments

This work was supported by the Polish National Science Centre (grant 2020/37/B/NZ2/03268). S.D.-H. was additionally supported by institutional funds from the Max Planck Society.

